# Characterizing a collective and dynamic component of chromatin immunoprecipitation enrichment profiles in yeast

**DOI:** 10.1101/004614

**Authors:** Lucas D. Ward, Junbai Wang, Harmen J. Bussemaker

**Affiliations:** Department of Biological Sciences, Columbia University, 1212 Amsterdam Ave., New York, NY 10027, USA; Center for Computational Biology and Bioinformatics, Columbia University, 1130 St. Nicholas Ave., New York, NY 10032, USA

**Keywords:** Transcription factors, chromatin immunoprecipitation, *Saccharomyces cerevisiae*

## Abstract

**Background:** Recent chromatin immunoprecipitation (ChIP) experiments in fly, mouse, and human have revealed the existence of high-occupancy target (HOT) regions or “hotspots” that show enrichment across many assayed DNA-binding proteins. Similar co-enrichment observed in yeast so far has been treated as artifactual, and has not been fully characterized.

**Results:** Here we reanalyze ChIP data from both array-based and sequencing-based experiments to show that in the yeast *S. cerevisiae*, the collective enrichment phenomenon is strongly associated with proximity to noncoding RNA genes and with nucleosome depletion. DNA sequence motifs that confer binding affinity for the proteins are largely absent from these hotspots, suggesting that protein-protein interactions play a prominent role. The hotspots are condition-specific, suggesting that they reflect a chromatin state or protein state, and are not a static feature of underlying sequence. Additionally, only a subset of all assayed factors is associated with these loci, suggesting that the co-enrichment cannot be simply explained by a chromatin state that is universally more prone to immunoprecipitation.

**Conclusions:** Together our results suggest that the co-enrichment patterns observed in yeast represent transcription factor co-occupancy. More generally, they make clear that great caution must be used when interpreting ChIP enrichment profiles for individual factors in isolation, as they will include factor-specific as well as collective contributions.

## BACKGROUND

In addition to mapping canonical transcription factor (TF) binding sites, chromatin immunoprecipitation (ChIP) experiments have revealed genomic loci at which many DNA-binding proteins display a signal of enrichment despite the absense of an *in vitro* binding site in the underlying DNA sequence. These regions have been alternatively called “TF colocalization hotspots” [1] and “high-occupancy target (HOT) regions” [2]. Their existence was first demonstrated in a study profiling seven *Drosophila melanogaster* TFs with diverse functions using the DamID method in cultured embryonic cells [1]. In that study, DNA at the hotspots was predicted to have affinity for three of the seven proteins (Gaf, Jra, and Max), but was bound by all seven. The hotspots were associated with increased expression at neighboring genes, suggesting that they are functionally relevant. Subsequent ChIP studies in whole embryos have confirmed that such hotspots are a general feature of the *Drosophila* [3–5] and the *C. elegans* [6] genomes. The TF colocalization phenomenon has also been observed in mammalian cells. An analysis of ChIP profiles for 13 TFs collected in mouse embryonic stem cells revealed extensive colocalization of these proteins along the genome [7]. Similarly, analysis of 89 sequence-specific TFs in a variety of human cell types [8] identified many HOT regions [2].

A number of mechanisms have been proposed to explain the observed co-enrichment across ChIP experiments. Chromatin loops could cross-link to multifunctional “transcription factories” or enhanceosomes [9]. Non-sequence-specific binding can also be driven by a locally permissive chromatin structure [3, 10]. The authors of the fly DamID study [1] argue against non-specific binding, because two non-endogenous proteins (mutant fly Bcd consisting of only a DNA-binding domain, and yeast Gal4p consisting of only a DNA-binding domain) do not localize to the hotspots, but rather to their predicted *in vitro* binding sites. Direct protein-protein interactions between the involved fly TFs have also not been observed, complicating any model involving a transcription factory. The authors of the mouse study [7], by contrast, suggest that the mouse hotspots represent enhanceosomes, due to their ability to drive transcription in a luciferase assay and their recruitment of the p300 coactivator. A feature shared by both organisms is that hotspots are associated with increased expression at neighboring genes, but are often located far from traditionally-defined proximal promoters.

The present study was motivated by the fact that, although extensive genome-wide *in vivo* protein binding data has been collected for the yeast *Saccharomyces cerevisiae* [11–13], no analogous colocalization of sequence-specific regulators has been reported for this organism. Significantly, however, in the large-scale compendia by Lee et al. [12] and Harbison et al. [11], the authors subtracted, for each probe separately, the mean across all arrays in order to account for biases in the immunoprecipitation reaction. This normalization procedure was certainly appropriate given the goal of these studies, namely, to determine the specific transcriptional target genes of each individual transcription factor. However, it would also have largely removed any true collective genomic enrichment pattern shared by many TFs. This insight motivated us to perform a detailed re-analysis of the original microarray data in a manner that omitted the probe-specific normalization step. This revealed that a collective pattern of ChIP enrichment also exists in yeast.

Unlike in higher eukaryotes, the collective enrichment patterns in yeast are not associated with sequence-predicted protein-DNA binding affinity for any of the TFs involved. Rather, sequence and functional analysis reveals that the most significant features of co-enriched probed regions are: (i) the extent of nucleosome depletion, (ii) expression of proximal genes, and (iii) the proximity to noncoding RNA genes, the majority of which encode tRNAs and snoRNAs. Additionally, the co-enrichment hotspots are occupied chiefly in rich-media (YPD) conditions, while, strikingly, the phenomenon is abrogated in the majority of environmental perturbation and stress conditions.

## RESULTS

### Quantifying collective ChIP enrichment in rich media conditions

First, we performed a detailed re-analysis of the raw ChIP-chip data from Lee et al. [12] and Harbison et al. [11], but *without* performing their normalization procedure across experiments (see Methods). To characterize the shared component of the ChIP profiles collected in rich media (YPD), we computed the median log_2_ fold-enrichment (MLFE) across 195 TFs as a measure of co-enrichment for each probe. The distribution of MLFE across probes was skewed heavily to the right (Figure 1), a shared enrichment profile that was evident in the authors’ original analysis but not fully characterized. The re-analyzed ChIP landscapes were also more correlated with each other than the normalized profiles from the original paper (Figure 2). We proceeded to investigate the location of the co-enrichment phenomenon relative to genomic features.

**Figure 1-.**
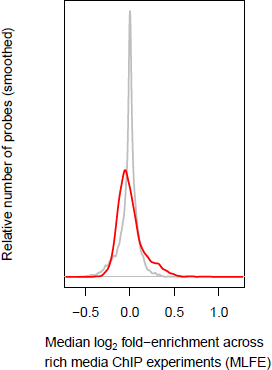
ChIP co-enrichment. Distribution of TF ChIP co-enrichment across probes. Co-enrichment is quantified as median log_2_ fold enrichment (MLFE) across all analyzed rich media experiments from Harbison et al. [11]. The distribution of the original normalized published data is in gray, and the distribution of the reanalyzed data is in red.

**Figure 2-.**
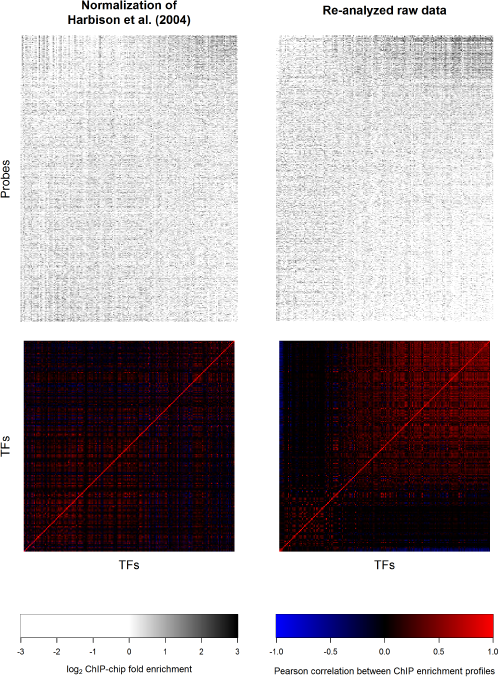
ChIP enrichment profiles from published and reanalyzed data. ChIP-chip enrichment profiles across all analyzed rich media experiments and correlations among them. An enrichment profile heatmap and correlatogram is shown for both the original normalized published data of Harbison et al. [11] and our reanalysis. TFs in all four matrices were sorted by their enrichment at ncRNA genes in the reanalyzed data; probes in the heatmaps were sorted by their median log_2_ fold enrichment (MLFE) in the reanalyzed data.

### Collective enrichment is strongly associated with noncoding-RNA genes

A first glance at the most highly co-enriched probed regions revealed a preponderance of telomeres and noncoding RNAs (ncRNAs) (Table 1). To systematically determine whether specific genomic features were associated with co-enrichment, we tested whether the distribution of MLFE for probes corresponding to each annotated genomic feature was different from that corresponding to the rest of the genome (Table 2). The most significantly co-enriched were the 514 probes corresponding to ncRNA genes (difference of median fold enrichment ΔMLFE = 0.27; *p* = 6.9 × 10^−161^, Student’s t-test; *p* < 2.2 × 10^−16^, Wilcoxon-Mann-Whitney test). The more specific ncRNA categories of tRNAs, snoRNAs, and snRNAs were all significantly co-enriched as well. There were not enough probes corresponding to rRNA genes to establish statistical significance.

**Table 1.**
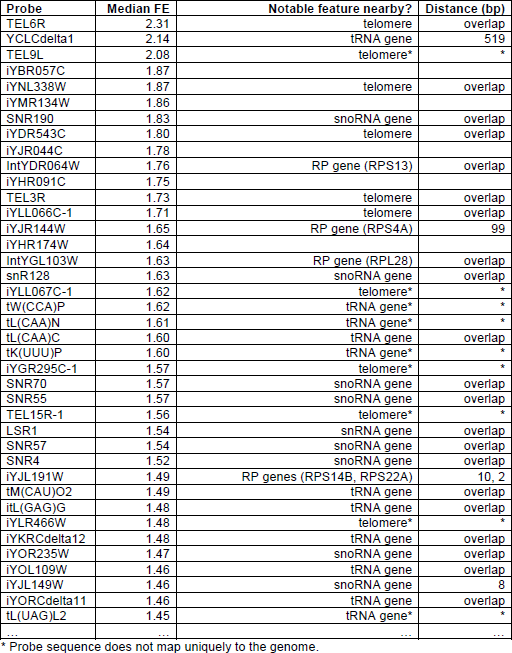
Probes with highest median ChIP-chip fold-enrichment (FE) across rich media experiments. List of probes with highest median ChIP-chip fold-enrichment (FE) across rich media experiments from Harbison et al. [11].

**Table 2.** Comparison of TF co-occupancy in each probe family vs. all other probes. Probes were mapped to a feature if there was any overlap between the probe and feature. For each probe, the co-occupancy was defined as the median log_2_ ChIP-chip fold enrichment (MFE) across all rich media experiments. For each feature, the probe family co-occupancy *ΔM̄* was defined as the difference in mean co-occupancy within each probe family and mean co-occupancy at all other probes. The *p*-value was determined using a t-test. Significant *p*-values are highlighted.

| Probe family/feature | Family size | Family TF co-enrichment | $-\log_{10}(p)$ |
| --- | --- | --- | --- |
| ncRNA | 514 | 0.27 | 160.16 |
| tRNA | 391 | 0.26 | 134.72 |
| snoRNA | 100 | 0.28 | 27.40 |
| repeat region | 369 | 0.07 | 8.55 |
| intron | 106 | 0.12 | 6.92 |
| telomere | 15 | 0.41 | 4.09 |
| snRNA | 13 | 0.24 | 3.45 |
| transposable element | 53 | 0.08 | 2.78 |
| ARS | 274 | -0.02 | 1.79 |
| transposable element gene | 6 | -0.05 | 0.93 |
| rRNA | 4 | -0.12 | 0.84 |
| pseudogene | 14 | 0.02 | 0.21 |
| centromere | 24 | -0.01 | 0.05 |

A subset of yeast tRNA genes have been demonstrated to colocalize to the nucleolus. We therefore asked whether TF co-enrichment is associated with nucleolar localization. We used the classification of yeast tRNA genes as nucleolar or non-nucleolar based on a three-dimension model of the yeast genome derived from chromatin conformation capture data by Duan et al. [14]. However, we found no significant difference in rich media MLFE between the two sets of genes (*t* = 0.67, *p* = 0.51). Therefore, nucleolar and centromeric tRNA genes seem to participate in the collective enrichment phenomenon to an equal degree.

### Evidence that collective enrichment is not due to technical artifact

Because telomeres and tRNA genes are associated with repetitive elements [15–16] in addition to having a high genomic copy number, we suspected that their consistently high enrichment across experiments could be an artifact of cross-hybridization [17–18]. To test for this, we inspected spot intensities and performed a more finely-grained classification of probes (Table 3; see Methods). We decided to exclude probes corresponding to telomeres or overlapping ncRNA genes by more than 25 bp from the remainder of our analysis (see Methods).

**Table 3.** Comparison of TF co-occupancy and absolute intensity among selected probe families and sub-families and different mapping criteria. TF co-enrichment *M* was defined for each probe as the median log_2_ fold enrichment across all rich media ChIP-chip experiments, and the family *ΔM̄* as the difference in mean *M* among probes in a family and all other probes. The *p*-value was calculated using a *t*-test. Similarly, the absolute intensity *A* for each probe in each experiment was defined as the mean (Lowess-normalized) intensity between the red and green channels; the median *A* was calculated across all experiments for each probe; and the family *ΔĀ* was reported as the difference in mean *A* among probes within a family and all other probes. Probe mapping and categories for comparison are as follows (see Methods for details of probe categorization): (A) Probes with high overlap vs. all other probes. (B) Probes with any overlap vs. all other probes. (C) Probes with low overlap or neighboring vs. non-neighboring probes (high overlap probes excluded from the analysis.) (D) Neighboring probes vs. non-neighboring probes (probes with any overlap excluded from the analysis.) Significant co-enrichment (*ΔM̄*) *p*-values are highlighted yellow; significant intensity (*ΔĀ*) *p*-values, which may signify cross-hybridization, are highlighted red.

**A**
| | n | $\Delta\bar{M}$ | $-\log_{10}(p)$ | $\Delta\bar{A}$ | $-\log_{10}(p)$ |
| --- | --- | --- | --- | --- | --- |
| ncRNA | 115 | 0.31 | 34.72 | -1.07 | 8.22 |
| tRNA | 33 | 0.37 | 16.37 | -1.99 | 9.59 |
| tRNA - spliced | 5 | 0.46 | 2.90 | -0.92 | 2.02 |
| tRNA - unspliced | 28 | 0.36 | 13.71 | -2.18 | 8.94 |
| tRNA - with Ty | 18 | 0.40 | 9.60 | -1.81 | 5.36 |
| tRNA - without Ty | 15 | 0.33 | 7.24 | -2.20 | 4.56 |
| snoRNA | 69 | 0.31 | 20.96 | -0.74 | 2.67 |
| snoRNA - C/D box | 36 | 0.37 | 12.60 | -1.76 | 5.85 |
| snoRNA - H/ACA box | 28 | 0.24 | 8.73 | 0.12 | 0.19 |
| snoRNA - unknown family | 5 | 0.25 | 1.34 | 1.81 | 2.43 |
| snoRNA - extragenic | 66 | 0.30 | 19.44 | -0.67 | 2.20 |
| snoRNA - intron-derived | 3 | 0.41 | 2.24 | -2.19 | 1.25 |
| snRNA | 6 | 0.35 | 2.28 | -1.18 | 0.81 |
| telomere | 12 | 0.49 | 4.04 | 2.19 | 2.67 |

**B**
| | n | $\Delta\bar{M}$ | $-\log_{10}(p)$ | $\Delta\bar{A}$ | $-\log_{10}(p)$ |
| --- | --- | --- | --- | --- | --- |
| ncRNA | 514 | 0.27 | 160.16 | -0.13 | 1.41 |
| tRNA | 391 | 0.26 | 134.72 | -0.04 | 0.24 |
| tRNA - spliced | 80 | 0.26 | 28.41 | 0.17 | 0.73 |
| tRNA - unspliced | 312 | 0.26 | 103.57 | -0.08 | 0.56 |
| tRNA - with Ty | 246 | 0.27 | 83.46 | -0.22 | 2.50 |
| tRNA - without Ty | 145 | 0.23 | 47.34 | 0.28 | 1.80 |
| snoRNA | 100 | 0.28 | 27.40 | -0.53 | 2.43 |
| snoRNA - C/D box | 58 | 0.31 | 16.93 | -1.06 | 4.33 |
| snoRNA - H/ACA box | 37 | 0.22 | 10.49 | 0 | 0 |
| snoRNA - unknown family | 5 | 0.25 | 1.34 | 1.81 | 2.43 |
| snoRNA - extragenic | 95 | 0.27 | 25.13 | -0.50 | 2.12 |
| snoRNA - intron-derived | 5 | 0.35 | 3.05 | -1.04 | 0.61 |
| snRNA | 13 | 0.24 | 3.45 | -0.53 | 0.63 |
| telomere | 15 | 0.41 | 4.09 | 2.23 | 3.74 |

C
| | n | $\Delta\bar{M}$ | $-\log_{10}(p)$ | $\Delta\bar{A}$ | $-\log_{10}(p)$ |
| --- | --- | --- | --- | --- | --- |
| ncRNA | 570 | 0.25 | 191.6 | 0.08 | 0.86 |
| tRNA | 500 | 0.25 | 172.62 | 0.07 | 0.66 |
| tRNA - spliced | 115 | 0.25 | 38.53 | 0.04 | 0.13 |
| tRNA - unspliced | 388 | 0.24 | 128.37 | 0.09 | 0.85 |
| tRNA - with Ty | 324 | 0.25 | 108.91 | -0.17 | 2.06 |
| tRNA - without Ty | 176 | 0.22 | 55.77 | 0.49 | 7.43 |
| snoRNA | 60 | 0.20 | 15.81 | 0.12 | 0.35 |
| snoRNA - C/D box | 47 | 0.21 | 13.71 | 0.23 | 0.76 |
| snoRNA - H/ACA box | 14 | 0.16 | 2.94 | -0.16 | 0.18 |
| snoRNA - unknown family | NA | NA | NA | NA | NA |
| snoRNA - extragenic | 58 | 0.20 | 14.87 | 0.10 | 0.27 |
| snoRNA - intron-derived | 2 | 0.27 | 1.10 | 0.68 | 1.40 |
| snRNA | 8 | 0.15 | 2.18 | -0.08 | 0.07 |
| telomere | 4 | 0.10 | 1.43 | 1.96 | 1.46 |

D
| | n | $\Delta\bar{M}$ | $-\log_{10}(p)$ | $\Delta\bar{A}$ | $-\log_{10}(p)$ |
| --- | --- | --- | --- | --- | --- |
| ncRNA | 171 | 0.24 | 56.57 | -0.06 | 0.27 |
| tRNA | 142 | 0.24 | 46.65 | -0.12 | 0.61 |
| tRNA - spliced | 40 | 0.25 | 11.91 | -0.34 | 1.13 |
| tRNA - unspliced | 104 | 0.23 | 34.12 | -0.02 | 0.05 |
| tRNA - with Ty | 96 | 0.24 | 30.84 | -0.32 | 1.98 |
| tRNA - without Ty | 46 | 0.22 | 14.14 | 0.28 | 0.80 |
| snoRNA | 29 | 0.20 | 8.17 | 0.29 | 0.75 |
| snoRNA - C/D box | 25 | 0.22 | 8.12 | 0.35 | 0.87 |
| snoRNA - H/ACA box | 5 | 0.15 | 0.90 | 0.23 | 0.16 |
| snoRNA - unknown family | NA | NA | NA | NA | NA |
| snoRNA - extragenic | 29 | 0.20 | 8.16 | 0.29 | 0.75 |
| snoRNA - intron-derived | NA | NA | NA | NA | NA |
| snRNA | NA | NA | NA | NA | NA |
| telomere | NA | NA | NA | NA | NA |

A plot of MLFE versus distance between the center of each probe and the center of the nearest ncRNA gene (Figure 3) shows a gradual and approximately exponential decay with increasing distance. The decay length is similar to a typical IP fragment length [19]. By contrast, cross-hybridization would appear as spikes as a function of genomic position with no such decay around peaks, as was discussed by Orian and colleagues [20]. We conclude that cross-hybridization is not responsible for the observed signal.

**Figure 3.**
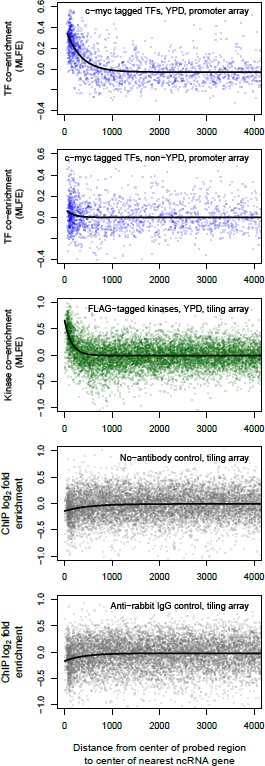
ChIP co-enrichment at ncRNA genes. TF co-enrichment, defined as the median log_2_ ChIP-chip fold enrichment (MLFE), as a function of distance to the nearest ncRNA gene. Plotted in black is a fit to *y = b_0_ + b_1_*e^(1/d)^. Top to bottom: (A) Co-enrichment across YPD experiments from Harbison et al. [11]: *b_0_* = −0.03, *b*_1_ = 0.44, *d* = 316.8; *p* for each parameter < 2 × 10^−16^; *r*^2^ = 0.27. (B) Co-enrichment across non-YPD experiments from Harbison et al. [11]: *b_0_* = −0.004, *b_1_* = 0.09, *d* = 154.3; *p* for each parameter > 0.04; r^2^ = 0.004. (C) Co-enrichment across YPD experiments from Pokholok et al. [27]: *b_0_* = −0.01, *b*_1_ = 0.67, *d* = 148.4; *p* for each parameter < 2 × 10^−16^; *r*^2^ = 0.04. (D) ChIP-chip log_2_ fold enrichment for no-antibody control from Pokholok et al. [23]; *b_0_* = −0.008, *b*_1_ = −0.13, *d* = 504.2; *p* for each parameter < 4.4 × 10^−10^; *r*^2^ = 0.004. (E) ChIP-chip log_2_ fold enrichment for anti-rabbit IgG control from Pokholok et al. [23]: *b_0_* = −0.02, *b*_1_ = −0.15, *d* = 417.1; *p* for each parameter < 4.9 × 10^−8^; *r*^2^ = 0.004.

Biases in IP efficiency and shearing based on chromatin state have been shown to be important in the interpretation of ChIP experiments [21–22]. To check whether such biases affected immunoprecipitation or hybridization efficiency of ncRNA genes, we inspected control experiments that used no antibody or a nonspecific antibody [23]. We observed a weak depletion of ncRNA genes in the mock IP samples relative to the whole-cell extract (no-antibody: ΔMLFE = −0.12; *p* = 2.9 × 10^−24^, t-test; rabbit IgG: ΔMLFE = −0.13; *p* = 1.9 × 10^−23^, t-test; Figure 3). These controls suggest that any immunoprecipitation bias at ncRNA genes would cause us to underestimate rather than overestimate the magnitude of the hotspot effect.

The ChIP-chip experiments that we re-analyzed for this study all relied on myc-tagged proteins. In humans, the c-Myc protein is localized to the nucleolus, raising the possibility that myc-tagged proteins in the ChIP experiment would be artificially biased towards tRNAs genes, some of which cluster in the nucleolus [24–26]. To rule out this possibility, we performed the same analysis on a set of ChIP-chip data that employed FLAG tagging rather than myc tagging, and high-density tiling probes [27].

The kinases assayed in this experiment again showed shared IP at ncRNA genes and exponential decay with increasing distance between the probed region and the ncRNA gene, and a comparable quantitative enrichment near ncRNA genes (ΔMLFE = 0.36; *p* = 2.1 × 10^−116^, t-test; Figure 3). Taken together, the above results make it unlikely that shared IP is dues to a tag-specific artifact.

### For most TFs, *in vitro* DNA binding specificity is a poor predictor of *in vivo* occupancy

The canonical view holds that the DNA-binding domain (DBD) of a TF is responsible for its recruitment to specific sequences in the genome. However, highly specific yet DBD-independent recruitment to sites of co-occupancy has been demonstrated using recombinant Bicoid protein in *Drosophila* [1]. The landscape of co-enrichment that we have characterized represents an independent contribution to the ChIP enrichment landscape of any given TF, which complements the sequence-specific targeting via its DBD. We were interested in contrasting these two predictors and quantifying the extent to which each of them contributes to the overall genomic enrichment profile for a TF. To this end, we calculated the Pearson correlation, across all probes, between the log_2_ fold enrichment (LFE) for each TF and (i) the median log_2_ fold-enrichment (MLFE) over all other TFs profiled in rich media, and (ii) the regional *in vitro* binding affinity predicted from DNA sequence using a position-specific affinity matrix for the TF from protein-binding microarray (PBM) data from Badis et al. [28] and Zhu et al. [29] (Figure 4). For almost all TFs, the correlation with MLFE is significant (mean value of *r* = 0.31), indicating that the co-enrichment signal contributes to their IP profile to a significant extent. A notable exception is Yap1p, whose LFE is significantly anticorrelated with the MLFE of all of other factors. For a smaller number of TFs, LFE correlates with predicted affinity, but always to a lesser extent than with MLFE (mean *r* = 0.04), with the exception of Abf1p.

**Figure 4.**
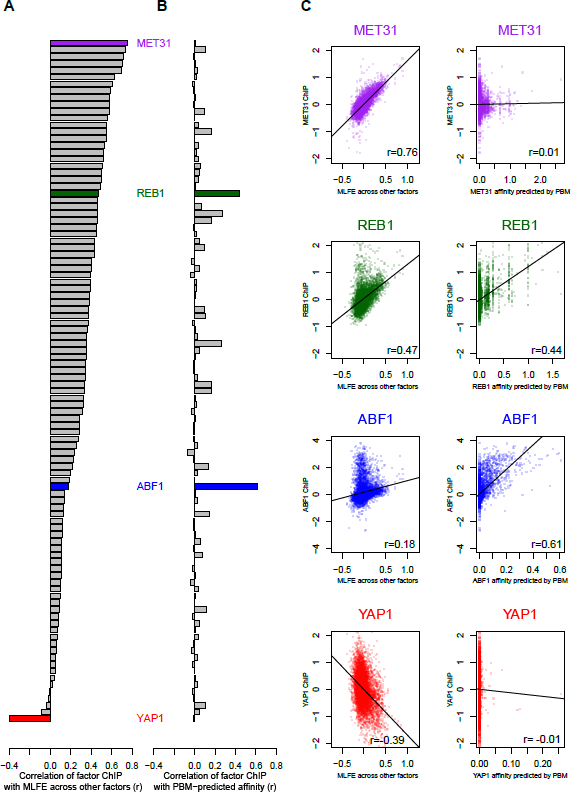
Correlation of ChIP enrichment for individual factors with co-enrichment and predicted affinity. Left to right: (A) Shared enrichment for each factor measured as the Pearson correlation between the TF’s genomewide enrichment landscape (in terms of log_2_ fold enrichment) and the median log_2_ fold enrichment (MLFE) across all other rich media ChIP-chip experiments. (B) Sequence-specific enrichment for each factor measured as the Pearson correlation between the TF’s genomewide enrichment and the predicted genomewide affinity for that TF from the PBM data of either Badis et al. [28] or Zhu et al. [29] (stronger correlation shown when TF is in both datasets). (C) Scatter plots showing the correlations described above (ChIP enrichment vs. co-enrichment and ChIP enrichment vs. affinity) for each of four factors: Met31p, Reb1p, Abf1p, and Yap1p.

### Co-enriched loci are associated with nucleosome depletion and high expression

To explore other relationships between genome function and TF co-enrichment, we looked for Gene Ontology (GO) categories of proximal genes (Table 4). For every GO category, we compared the distribution of MLFE within probes corresponding to promoters of genes in that category with the rest of the probes. The most enriched protein functions are for translation (translational elongation, t = 13.8, *p* = 7.7 × 10^−43^; cytoplasmic translation, t = 13.8, *p* = 1.1 × 10^−42^) and accordingly, ribosomal proteins as a whole are strongly enriched (t = 10.4, *p* = 3.1 × 10^−25^). Because ribosomal protein (RP) promoters are known to be particularly active [30], we were interested in whether expression globally correlates with co-enrichment, and found that it does (Pearson *r* = 0.17, *p* = 1.1 × 10^−40^; Figure 5A). We also found that co-enrichment is even more strongly anticorrelated with nucleosome occupancy (Pearson *r* = –0.31, *p* = 1.3 × 10^−122^; Figure 5B).

**Figure 5.**
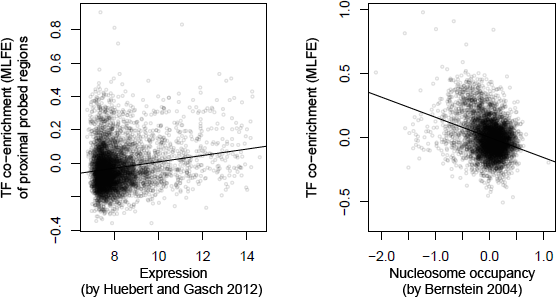
Correlation of TF co-enrichment with gene expression and nucleosome occupancy. (L) Scatter plot of TF co-enrichment vs. gene expression in YPD from Huebert and Gasch [48]. Each point is the expression level for a gene and the co-enrichment (MLFE) of neighboring regions; expression values are log_2_ of quantile normalized intensity values. Plotted as a black line is a fit of all the data to a linear model (*r* = 0.17). (R) Scatter plot of TF co-enrichment vs. nucleosome occupancy by nucleosome ChIP from Bernstein et al. [31]. Each point is a probed region assayed both by Harbison et al. and Bernstein et al. Plotted as a line is a fit of all the data to a linear model (*r* = −0.31).

**Table 4.** Gene Ontology (GO) enrichment analysis of genes by level of TF co-enrichment at neighboring probes.

| **GO Category** | **p-value** | **t** |
| --- | --- | --- |
| 6414\|P\|translational_elongation | 7.74E-43 | 13.84 |
| 2181\|P\|cytoplasmic_translation | 1.11E-42 | 13.81 |
| 30533\|F\|triplet_codon-amino_acid_adaptor_activity | 2.65E-38 | 13.04 |
| 3735\|F\|structural_constituent_of_ribosome | 9.10E-32 | 11.80 |
| 22625\|C\|cytosolic_large_ribosomal_subunit | 5.13E-31 | 11.65 |
| 5840\|C\|ribosome | 3.10E-25 | 10.43 |
| 6412\|P\|translation | 1.04E-23 | 10.08 |
| 22627\|C\|cytosolic_small_ribosomal_subunit | 2.43E-21 | 9.52 |
| 943\|C\|retrotransposon_nucleocapsid | 4.54E-20 | 9.21 |
| 30529\|C\|ribonucleoprotein_complex | 6.04E-20 | 9.18 |
| 32197\|P\|transposition,_RNA-mediated | 1.49E-19 | 9.08 |
| 5829\|C\|cytosol | 9.69E-17 | 8.34 |
| 5622\|C\|intracellular | 9.59E-13 | 7.15 |
| 15074\|P\|DNA_integration | 1.66E-09 | 6.04 |
| 46797\|P\|viral_procapsid_maturation | 1.66E-09 | 6.04 |
| 3964\|F\|RNA-directed_DNA_polymerase_activity | 8.01E-09 | 5.78 |
| 3723\|F\|RNA_binding | 1.55E-08 | 5.66 |
| 4523\|F\|RNA-DNA_hybrid_ribonuclease_activity | 3.14E-08 | 5.54 |
| 4540\|F\|ribonuclease_activity | 5.65E-08 | 5.44 |

We were also interested in whether the TF co-enrichment profile was correlated with affinity for TFs. We calculated the predicted affinity of each probe for a compendium of TFs. Among TF affinities predicted from protein binding microarray (PBM) data, only affinity for Rsc30p, Rsc3p, and Rap1p correlated with MLFE (Pearson *r* = 0.07, *r* = 0.06, and *r* = 0.06, respectively). Binding by these factors has previously been shown to drive nucleosome depletion at RP promoters [28, 31], consistent with the correlation with nucleosome depletion described above.

### Collective enrichment at ncRNA genes is largely eliminated in perturbed conditions

So far, our analysis has been restricted to rich media (YPD) conditions, providing a uniform chromatin context for comparison across factors. Examining ncRNA loci in experimental perturbation (“stress”) conditions reveals dramatically reduced co-enrichment (Table 5 and Figure 3). Using the median TF enrichment across all non-YPD conditions, the elevation in co-enrichment at ncRNA genes drops from 0.25 to 0.03. To further investigate this general observation by focusing on ChIP enrichment of individual TFs in their rich media and stress conditions. For each particular stress-TF combination (i.e., each experiment), we calculated the enrichment at ncRNA genes relative to all other probes (Figure 6). As expected from our pooled analysis, in the majority of stress conditions the enrichment at ncRNA genes is greatly reduced. For two TFs, viz. Kss1p and Gal4p, ncRNA genes are preferentially ChIP enriched in YPD, while in stress the enrichment at ncRNA genes is lower than elsewhere in the genome. Kss1p shows a negative relative occupancy of ncRNA genes in alpha mating factor and 1-butanol conditions. Gal4p shows decreased preferential enrichment at ncRNA genes in galactose and avoidance of these loci in raffinose.

**Figure 6.**
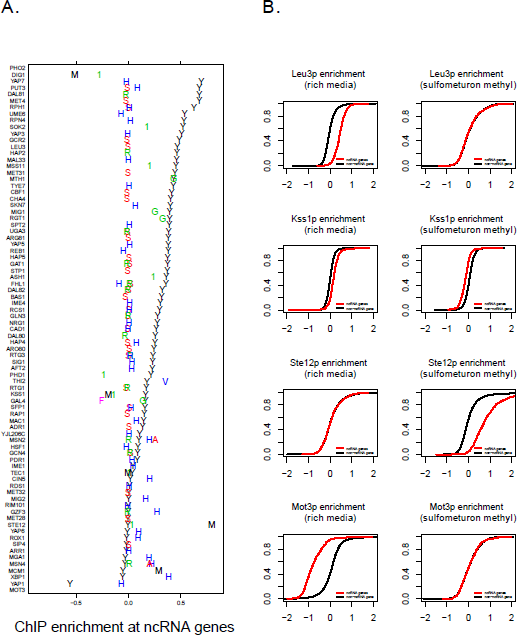
Condition specificity of co-enrichment at ncRNA genes. (A) Each row is a TF, and experimental conditions for that TF are plotted on the same row with letters indicating the condition. Conditions are: “Y”, rich media; “S”, sulfometuron methyl; “R”, rapamycin; “H”, hydrogen peroxide; “1”, 1-butanol; “A”, succinic acid; “G”, galactose; “V”, vitamin deprived medium; “M”, alpha mating factor; “F”, raffinose; and “P”, phosphate deprived medium. ChIP enrichment at ncRNA genes is expressed as the difference between the mean log_2_ fold enrichment of ncRNA gene probes and the mean log_2_ fold enrichment of all other probes. (B) Leu3p, Ste12p, and Mot3p enrichment at ncRNA genes in rich media vs. sulfometuron methyl treatment. For each factor and condition, an empirical cumulative distribution function is shown contrasting the distribution in log_2_ fold enrichment (FE) for ncRNA gene probes and all other probes.

**Table 5.** Comparison of TF co-enrichment for ncRNA families in rich media and stress conditions. Criterion for probe mapping is the same as in Table 3C: Probes with low overlap or neighboring vs. non-neighboring probes (high overlap probes excluded from the analysis.) Significant *p-*values are highlighted in yellow.

| | n | $\Delta\bar{M}$ (YPD) | $-\log_{10}(p)$ | $\Delta\bar{M}$ (stress) | $-\log_{10}(p)$ |
| --- | --- | --- | --- | --- | --- |
| ncRNA | 570 | 0.25 | 191.6 | 0.03 | 5.92 |
| tRNA | 500 | 0.25 | 172.62 | 0.03 | 5.51 |
| tRNA - spliced | 115 | 0.25 | 38.53 | 0.05 | 3.18 |
| tRNA - unspliced | 388 | 0.24 | 128.37 | 0.03 | 3.07 |
| tRNA - with Ty | 324 | 0.25 | 108.91 | 0.03 | 2.73 |
| tRNA - without Ty | 176 | 0.22 | 55.77 | 0.03 | 2.92 |
| snoRNA | 60 | 0.2 | 15.81 | 0.01 | 0.15 |
| snoRNA - C/D box | 47 | 0.21 | 13.71 | 0.01 | 0.11 |
| snoRNA - H/ACA box | 14 | 0.16 | 2.94 | 0.01 | 0.09 |
| snoRNA - extragenic | 58 | 0.2 | 14.87 | 0.01 | 0.17 |
| snoRNA - intron-derived | 2 | 0.27 | 1.1 | -0.03 | 0.19 |
| snRNA | 8 | 0.15 | 2.18 | -0.08 | 1.15 |

Interestingly, those TFs that do not participate in ChIP co-enrichment at ncRNA genes as strongly in rich media conditions are more likely to be ChIP-enriched at ncRNA genes in other conditions. The most notable example of this is Ste12p, which is enriched at ncRNA genes upon exposure to alpha mating factor, but not in the absence of alpha factor or in the presence of 1-butanol. Dig1p, which is also associated with the mating response, behaves differently: it is not enriched at ncRNA genes in rich media, and is also not enriched at them in the presence of alpha mating factor and 1-butanol. Finally, among the other TFs that exhibit ncRNA depletion in rich media, Mot3p shows a loss of this depletion in the presence of hydrogen peroxide or sulfometuron methyl. The fact that enrichment at ncRNA genes is both factor and condition specific supports that the ChIP co-enrichment is not solely determined by the chromatin state at the co-enriched loci, and is dependent on the identity and activity of the binding proteins.

### Co-enrichment during oxidative stress is reduced, not moved to other loci

To directly compare co-enrichment between YPD and perturbed conditions, we looked at the hydrogen peroxide condition, which has the highest number of factors assayed in common with YPD. We then calculated MLFE in each condition using only the subset of factors that was assayed in both, and performed GO analysis (Table 6) and expression correlation analysis (Figure 7). Analyzing this subset, we again found the strongest co-enrichment at promoters of ribosome-associated genes, in both YPD and hydrogen peroxide conditions (Table 6). However, the enrichment was greatly reduced during oxidative stress, to the extent that only one GO category (small ribosomal subunit; see highlighted row) in the H_2_O_2_ condition showed an enrichment surpassing a threshold of p < (0.05/748 categories). In addition, the correlation between co-enrichment and expression is much weaker during oxidative stress (YPD r = 0.17, slope = 0.02 ± 0.002, p = 1.37 × 10^−35^; H_2_O_2_ r = 0.05, slope = .005 ± 0.001, p = 3.1 × 10^−4^).

**Figure 7.**
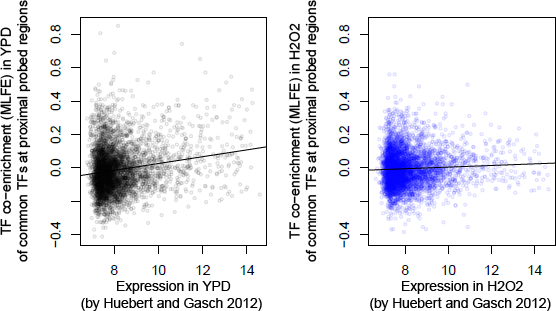
Correlation of TF co-enrichment with gene expression in different conditions. Scatter plots of TF co-enrichment vs. gene expression in YPD from Huebert and Gasch [48]. In each case, the co-enrichment (MLFE) is defined by using only data from TFs assayed in both YPD and H_2_O_2_. Each point is the expression level for a gene and the co-enrichment (MLFE) of neighboring regions; expression values are log_2_ of quantile normalized intensity values. (L) YPD, (R) H_2_O_2_.

**Table 6.** Gene Ontology (GO) enrichment analysis of genes by level of TF co-enrichment in both YPD and H_2_O_2_ conditions at neighboring probes. TFs in this analysis were restricted to the subset shared between YPD and H_2_O_2_ conditions, and the top GO enrichments are shown for YPD. The highlighted row is the only category that is significant in H_2_O_2_ after Bonferroni correction. Expression values were obtained from Huebert and Gasch (2012) as described in Methods.

|  | YPD p | YPD t | H<sub>2</sub>O<sub>2</sub> p | H<sub>2</sub>O<sub>2</sub> t |
| --- | --- | --- | --- | --- |
| 2181\|P\|cytoplasmic_translation | 2.12E-38 | 13.06 | 2.65E-03 | 3.01 |
| 3735\|F\|structural_constituent_of_ribosome | 3.14E-31 | 11.69 | 1.03E-02 | 2.57 |
| 22625\|C\|cytosolic_large_ribosomal_subunit | 6.66E-30 | 11.43 | 4.30E-02 | 2.02 |
| 5840\|C\|ribosome | 2.91E-27 | 10.87 | 1.20E-03 | 3.24 |
| 6414\|P\|translational_elongation | 4.76E-24 | 10.16 | 1.45E-01 | 1.46 |
| 6412\|P\|translation | 1.22E-23 | 10.07 | 5.46E-02 | 1.92 |
| 22627\|C\|cytosolic_small_ribosomal_subunit | 4.76E-21 | 9.45 | 1.03E-03 | 3.28 |
| 30533\|F\|triplet_codon-amino_acid_adaptor_activity | 2.24E-19 | 9.03 | 1.24E-01 | 1.54 |
| 30529\|C\|ribonucleoprotein_complex | 9.82E-19 | 8.87 | 1.47E-01 | 1.45 |
| 5622\|C\|intracellular | 5.83E-15 | 7.83 | 1.35E-01 | 1.49 |
| 943\|C\|retrotransposon_nucleocapsid | 3.35E-13 | 7.30 | 9.80E-03 | 2.58 |
| 32197\|P\|transposition,_RNA-mediated | 4.83E-13 | 7.25 | 1.21E-02 | 2.51 |
| 5829\|C\|cytosol | 4.27E-10 | 6.26 | 3.19E-02 | 2.15 |
| 6407\|P\|rRNA_export_from_nucleus | 8.68E-09 | 5.76 | 8.69E-03 | 2.62 |
| 15935\|C\|small_ribosomal_subunit | 2.54E-07 | 5.16 | 9.64E-05 | 3.90 |
| 15074\|P\|DNA_integration | 3.34E-06 | 4.65 | 1.62E-01 | 1.40 |
| 46797\|P\|viral_procapsid_maturation | 3.34E-06 | 4.65 | 1.62E-01 | 1.40 |
| 4540\|F\|ribonuclease_activity | 9.62E-06 | 4.43 | 2.95E-01 | 1.05 |
| 3964\|F\|RNA-directed_DNA_polymerase_activity | 1.14E-05 | 4.39 | 2.56E-01 | 1.14 |

### Validation by ChIP-Seq

For validation purposes, we compared three Ste12p ChIP-Seq datasets, one of which was performed in pseudohyphal conditions and two in exposure to alpha mating factor [32–33]. Both showed enrichment near ncRNA genes, although the magnitude was greater during exposure to alpha mating factor, consistent with the experiments of Harbison et al. [11] (Figure 8). These data further support that the hotspot effect is not an artifact of microarray technology.

**Figure 8.**
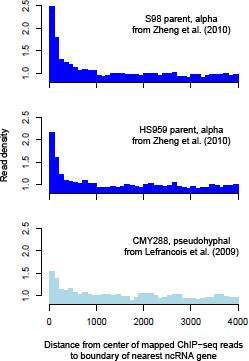
Validation using ChIP-Seq data. Density of Ste12p ChIP-seq reads relative to the genome-wide coverage for the two parents tested under exposure to alpha mating factor in Zheng et al [33] and the strain tested under pseudohyphal growth conditions in Lefrancois et al. [32].

## DISCUSSION

### Other evidence for TF colocalization in the yeast literature

Our reanalysis of the ChIP-chip compendia of Lee *et al.* [12] and Harbison *et al.* [11] has revealed co-enrichment of yeast TFs at ncRNA genes. In a more recent study, Venters and colleagues used low-density tiling microarrays to assay the occupancy of a broader range of factors [13]. Because of differences in probe design, their occupancy data are not directly comparable to those of Lee et al. [12] and Harbison et al. [11], and are not suited to the interrogation of transcribed regions; however, the authors noted a surprising association of Pol II-associated factors with tRNA promoters. Two recent studies in yeast have recognized non-canonical binding in light of the known biological roles of TFs. Fan and Struhl [34] found condition-specific Mediator binding over many gene bodies, rather than upstream promoter regions where it is known to act; they argued based on the low enrichment and reproducibility that these targets represent indirect binding due to chromatin state. Teytelman and colleagues [35], motivated by finding components of the Sir silencing complex at actively-transcribed regions, found that exogenously expressed GFP also immunoprecipitated with these regions in a condition-specific manner.

### Possible mechanisms underlying dynamic co-enrichment at ncRNA genes

Genomic recruitment of transcription factors is usually conceptualized as binding of the DNA-binding domain of the protein to high-affinity consensus sequences in the DNA, contingent on the local accessibility of the DNA. Our finding that many studied yeast transcription factors preferentially immunoprecipitate with nucleosome-depleted DNA is consistent with previous observations that TFs will nonspecifically bind to naked DNA at a low level [36]. Within the nucleus, nucleosome-depleted regions may most closely resemble naked DNA *in vitro*, in which case they ought to display a higher level of nonspecific binding relative to nucleosome-occupied and heterochromatic regions. However, we have shown here that the hotspot phenomenon can only be partly explained in terms of chromatin accessibility, because even when using the same antibody, the ChIP enrichment at hotspots depends on which TF carries the affinity tag. This is consistent with the recent observation in fly Kc cells [37] and in cultured human cells [38] that the optimal chromatin context – i.e., the chromatin type for which the highest degree of occupancy is observed at a given level of sequence-predicted DNA binding affinity – is different for each TF, and that none of the chromatin states is globally permissive.

Both the ChIP and DamID method can detect TFs that are near DNA but not necessarily contacting it. Consequently, the observed co-enrichment signal could be due to the proximity of probed regions to the TFs rather than due to direct interactions with them. Indeed, for individual yeast TFs, indirect interactions have been proposed in order to account for the poor correlation between *in vitro* sequence specificity as measured by protein binding microarrays (PBMs) and *in vivo* occupancy as measured by ChIP-chip [39]. Fly and mouse hotspots have been hypothesized to reflect both direct interactions mediated by the DNA-binding domain of certain TFs and indirect, protein-protein interactions involving the other co-enriched TFs [1, 7]. Our sequence analysis does not provide any evidence of direct sequence-specific interactions with TFs. Nucleosome depletion and proximity to ncRNA genes both predict co-enrichment significantly better than local regional binding affinity predicted from DNA sequences using either known binding specificities or *de novo* motif discovery. The co-enrichment could also be the result of competitive binding by different TFs in different subsets of cells and at different times, as has been suggested by recent work [40–41].

Several lines of cytological evidence from mammalian cells suggest that transcription by polymerase II occurs at nuclear foci comprising many polymerase molecules and transcription factors, termed “transcription factories” [9]. If such factories exist in yeast, it is conceivable that nucleosome-free regions and ncRNA genes – which are associated with high levels of transcription (by polymerase II and I/III, respectively) – are in close proximity to multiple TFs as a result of transcription factories. Indeed, it was recently discovered that Pol II-associated transcription factors tightly associate with Pol III-transcribed genes in human cells [42].

## CONCLUSIONS

Our results show that the median enrichment across all TFs is far more predictive of the ChIP landscape of a typical individual yeast TF than DNA sequence is. This agrees with a recent study of the interaction between chromatin accessibility and sequence specificity [10]. While the normalized enrichment data of the original yeast ChIP-chip compendia [11–12] have proven immensely valuable for understanding and modeling regulatory networks, any other ChIP experiment not subjected to the same normalization will display both sequence-specific as well as hotspot targeting. As genomic protein occupancy mapping technology increases in resolution and sensitivity, understanding the structure, origin, and possible function of co-enrichment hotspots will become increasingly important to interpreting the data they generate.

## METHODS

### Processing of raw ChIP-chip data

The original raw ChIP-chip data [11–12] were obtained from ArrayExpress (http://www.ebi.ac.uk/microarray-as/ae/) using accession numbers E-WMIT-1 and E-WMIT-10, respectively. Protocol information for each array (which dye was IP vs. WCE, experimental conditions, etc.) was extracted from the files E-WMIT-1.sdrf.txt and E-WMIT-10.sdrf.txt, available in the directory http://www.ebi.ac.uk/microarray-as/ae/download/. Raw intensity information was downloaded from the tab-delimited text files in E-WMIT-1.raw.zip and E-WMIT-10.raw.zip available in the FTP directory specified within the aforementioned text files. The column headers in all of these text files were found to be corrupted. Therefore, they were split between nine different formats. Each format was manually curated to locate the correct median foreground and background red and green intensity columns, using the presence of a background-subtracted log ratio column as a validation. Four of the experimental conditions had array data in the database that also had corrupted rows, where the number of columns was not consistent throughout the whole file; data associated with these conditions (Dal81p sulfometuron methyl, Arg80p sulfometuron methyl, Mac1p hydrogen peroxide, and Ime1 hydrogen peroxide) were discarded. Raw intensities were loaded into *R* and Loess normalization was performed on each array (to account for dye-specific response functions) using the normalizeWithinArrays function of the *limma* package [43], resulting in an M (relative intensity) and A (absolute intensity) value for each spot on each array. A number of the arrays were found to have very low variance in their log ratios; arrays with a variance in M after Loess normalization less than 0.05 were discarded. Four summary values were calculated for each probe: a median log ratio (M) and intensity (A) signal across all rich-media (YPD) arrays, and a median log ratio (M) and intensity (A) across all stress arrays. Additionally, for every experimental condition for which multiple replicates were available, a median M and A value across replicates was calculated. The same processing was applied to ArrayExpress data from assaying rabbit IgG control, no-antibody control, and kinase occupancy by tiling array [23, 27], which we used for validation.

### Genome annotation

The genomic coordinates of probes were mapped to the chromosome sequences contained in the GFF-formatted sequence and annotation available from the Saccharomyces Genome Database (SGD) [44], dated 21 April 2007, and the distance from each probe to the nearest annotated genomic feature of each type was calculated. More specifically, both a gap (defined as zero if overlapping, and otherwise the distance between the edge of a probe and the edge of a feature) and an overlap were calculated. The GFF file was further parsed to divide tRNAs into spliced vs. intronless and *Ty*-flanked vs. *Ty*-absent tRNAs, and to divide snoRNAs into H/ACA-box vs. C/D-box and intron-derived vs. extragenic snoRNAs. The array design includes both probes that are centered on tRNAs, and probes that only overlap partially with tRNAs. For each category of genomic feature, we defined the probes that were centered on the feature (“high overlap” > 25 bp), those with a partial overlap (“low overlap” ≤ 25 bp), those that were neighboring (“neighbors,” no overlap, gap between 1 and 100 bp), and all other probes.

### Annotation-specific inspection of intensities to test for cross-hybridization

In order to test for cross-hybridization, we inspected median intensities and performed *t*-tests for probes corresponding to each class and sub-class of features defined above (Table 3). While probes corresponding to telomeres had higher median log_2_ fold enrichment (MLFE; ΔMLFE = 0.41, *p* = 8.1 × 10^−5^), they also had higher median intensities (*ΔĀ* = 2.23, *p* = 1.8 × 10^−4^). Therefore, we excluded them from our analysis. Additionally, many families of ncRNA probes had *lower* median intensities, presumably due to their relatively short length. Using a conservative criterion for classification (Table 3D), discarding probes overlapping ncRNA genes and only considering neighboring probes, still results in the co-occupancy effect among the neighboring probes, suggesting that neither the high copy number of tRNAs nor of their associated *Ty* elements are responsible for the co-occupancy effect. We settled on a criterion that excludes any probes showing any overlap with telomeres or high overlap with ncRNA genes from the remainder of our analyses, but we did include probes neighboring ncRNA genes and those with a low overlap with ncRNA genes in our definition of ncRNA gene probes (the criterion used in Table 3C). A similar criterion was employed in a RNA polymerase III location study using a similar ChIP-chip array design [45].

### Comparison of occupancy at annotated targets and at ncRNA genes

Annotated targets for each TF were defined as probes that overlapped or were neighboring (within 100 bp of) regions reported by MacIsaac et al. [46] within their *p*-value threshold of 0.005. After discarding probes that were annotated both as ncRNA probes (according to the criterion described above) and as TF targets, we compared the mean log_2_ fold enrichment among ncRNA probes and among annotated targets with that of all other probes. A significant difference in means was defined as a *t*-test passing a *p*-value threshold of 0.05, Bonferroni corrected for the number of tests.

### Correlation with sequence-predicted binding affinity, nucleosome affinity, and gene expression

The affinity of each probed region for TFs was calculated using two published libraries of protein binding microarray (PBM)-derived position weight matrices (PWMs) [28–29]. The PWMs were converted to position-specific affinity matrices (PSAMs) and probe-TF affinities were calculated using the *AffinityProfile* utility in the *MatrixREDUCE* package as described previously [47]. The Pearson correlation between predicted affinity and MFE was then calculated. Nucleosome occupancy measurements by ChIP-chip were obtained from Bernstein et al. [31]. For each probed region, the median log ratio across all assayed histone subunits was used. The Pearson correlation between predicted affinity and nucleosome occupancy was then calculated. Gene expression data from both YPD and the 30 minutes treatment with 0.4mM concentration H_2_O_2_ condition were obtained from [48], and probes were assigned to genes using S. cerevisiae chromosomal features (Genome Version R64-1-1) annotated in Saccharomyces genome database (SGD). In cases of divergent promoters, value was assigned to both genes. Probe intensities were quantile normalized using MATLAB bioinformatics toolbox.

### Gene Ontology analysis

Functional enrichment of probes by Gene Ontology (GO) categories [49] was determined using a MATLAB implementation of the T-profiler algorithm [50]. The GO annotation was downloaded from SGD (Gene Ontology Consortium Validation Date: 01/25/2014).

### Condition specific analyses

Condition specificity was calculated as follows: For each YPD experiment, we calculated the Pearson correlation between the TF’s occupancy and the median occupancy across all other rich media TF experiments, and also the correlation between its occupancy and its predicted affinity as predicted from PBM data. TFs for which no PBM-derived matrix was available were excluded. In cases for which two matrices were available – from both Badis et al. [28] and Zhu et al. [29] – the one with the best correlation to the ChIP occupancy was used.

### ChIP-seq analysis

ChIP-seq data from Lefrancois et al. [32] and Zheng et al. [33] were downloaded from Gene Expression Omnibus (http://www.ncbi.nlm.nih.gov/geo). These data include mapped peaks, but not genome-wide mapping of reads; therefore, read alignment results from ELAND were downloaded and processed using MACS [50] as described by the authors in order to obtain a genome-wide landscape of binding, in 10-bp bins. Distances from these bins to ncRNA genes were measured using the SGD genome annotation described above and BEDTools [51].

## ABBREVIATIONS

ChIP: chromatin immunoprecipitation
DamID: DNA adenine methyltransferase identification
DBD: DNA-binding domain
IgG: immunoglobulin G
LFE: log_2_ fold enrichment
HOT: high-occupancy target
MLFE: median log_2_ fold enrichment
ncRNA: noncoding RNA
PBM: protein-binding microarray
RP: ribosomal protein
TF: transcription factor
YPD: yeast extract peptone dextrose

## AUTHOR CONTRIBUTIONS

The authors declare no conflicts of interest. LDW, JW, and HJB designed and performed research; LDW and HJB wrote the paper.

## ACKNOWLEDGEMENTS

We thank Bas van Steensel, Helen Causton, and members of the Bussemaker, Botstein, and Kellis labs for helpful discussions. This work was supported by Human Frontier Science Program (HFSP) grant RGP56/2003, National Institutes of Health (NIH) grants R01HG003008, U54CA121852, and T32GM082797, and P50GM071508, as well as a John Simon Guggenheim Foundation Fellowship to H.J.B.

